# Deep learning inference of cell type-specific gene expression from breast tumor histopathology

**DOI:** 10.1101/2025.05.04.652089

**Authors:** Andrew T. Wang, Saugato R. Dhruba, Kun Wang, Emma M. Campagnolo, Eldad D. Shulman, Eytan Ruppin

## Abstract

Cell type-specific gene expression from single-cell RNA sequencing (RNA-seq) is valuable for breast cancer precision oncology but available cohorts are still limited due to its high cost. Deconvolution methods infer cell type-specific expression from bulk RNA-seq at a lower cost, yet expenses and processing time of bulk RNA-seq are also non-negligible and limit their application too. To address these limitations, we developed *SLIDE-EX* (SLide-based Inference of DEconvolved gene EXpression), a deep-learning tool that predicts cell type-specific gene expression and abundances directly from routine breast cancer histopathology whole slide images (WSIs), using deconvolved bulk RNA-seq data as training labels. Trained on the TCGA-breast cohort and tested in cross validation and on an independent cohort of 160 cases, *SLIDE-EX* robustly infers the expression of thousands of genes across 9 distinct cell types, performing best for cancer associated fibroblasts and cancer cells. The abundance of these two cell types could also be robustly predicted, together with that of myeloid cells. The robustly predicted genes reflect key biological functions of their respective cell types. From a translational perspective, the inferred cell type specific expression profiles predict chemotherapy response more accurately than models based on direct prediction from the slides or from the inferred bulk expression in two independent cohorts. Going forward, *SLIDE-EX* is a generic approach that opens up possibilities for rapid, cost-effective cell type-specific gene expression inference in potentially any cancer type, further democratizing the characterization of the tumor microenvironment.

## Introduction

Breast cancer remains the most common cancer among women worldwide, with over 2 million new cases diagnosed annually^1^. This heterogeneous disease is classified into multiple subtypes, each with distinct biological features and clinical outcomes^2^. Despite advances in early detection and treatment, accurately predicting patient prognosis and response to therapy remains challenging, highlighting the need for more precise diagnostic and prognostic tools. In recent years, transcriptomic signatures have emerged as powerful tools in breast cancer precision oncology. These signatures provide crucial insights into tumor biology, helping to guide treatment decisions and identify patients who may benefit from specific therapies^3^. For example, the PAM50 gene expression assay measures the expression of 50 classifier genes to categorize breast cancer into molecular subtypes and predict clinical outcomes^4^. The Oncotype DX, one of the most widely used gene expression assay in clinical practice, generates a recurrence score that serves as a prognostic biomarker and predicts the benefits from chemotherapy in patients meeting specific diagnostic criteria^5–7^. Moreover, gene expression patterns have shown promise in identifying novel biomarkers and predicting response to both standard chemotherapy and targeted treatments^8,3^. These advances suggest that continued development and validation of gene expression signatures hold promise for improving patient stratification and treatment selection in clinical practice.

Beyond tumor bulk gene expression, the tumor microenvironment (TME) is emerging as crucial for patient stratification and treatment response prediction. It has been shown that the clustering and morphology of tumor-infiltrating lymphocytes, as well as the density and clustering of cancer-associated fibroblasts, are prognostic in invasive breast cancer^9^. Effective tools to study the various cellular states of different cell types include spatial transcriptomics and single-cell RNA sequencing, but their use remains limited due to high cost and scarcity^10^. A more cost-effective approach to identifying clinically relevant cellular states is extracting cell type-specific expression from bulk tumor gene expression samples through computational deconvolution methods, and several algorithms have been developed for this purpose^11–13^. Most deconvolution algorithms utilize reference single-cell signatures. CIBERSORTx applies support vector regression, using these signatures as features to infer cell type proportions and gene expression^11^. MuSiC incorporates cross-sample variation to refine predictions, adjusting references to better fit bulk data^12^. CODEFACS, used in this study, enhances deconvolution through iterative optimization, predicting more genes with higher accuracy^13^. However, the potential clinical application of these methods remains limited by their reliance on bulk RNA-seq assays, which are costly and have long turnaround times, thereby constraining the use of gene expression-based biomarkers for TME analysis and treatment response prediction.

Recent advances in digital histopathology have enabled the extraction of clinically significant information from tumor slides using machine learning and artificial intelligence techniques, leveraging rapid progress in deep learning-based computer vision^14^. The application of deep learning to whole-slide images (WSIs) of tissue stained with hematoxylin and eosin (H&E) has already demonstrated potential in detecting features beyond human perception, including genetic mutations^15^, bulk mRNA expression^16^, and methylation patterns^17^. H&E slides are widely available and routine during cancer screening and diagnosis, making them ideal for greater accessibility of predictive cancer tools. Previous work to predict transcriptomics from H&E images has centered on bulk expression^16,18–20^, and this framework has been successfully used to predict cancer treatment response from inferred bulk expression intermediates in independent, retrospective clinical trial datasets^16^. However, prediction of cell type-specific gene expression from H&E slides remains unexplored. Therefore, given the importance of transcriptomic signatures and expression in breast cancer treatment strategies, as well as recent developments in deconvolution methods and AI-driven biomarker prediction, **we aim to investigate two main research questions: (1) To what extent can we predict cell type-specific data—both gene expression and cell abundance—from slide images? (2) Given a cell type-specific predictor, to what extent can we use these predictions to infer clinically relevant phenotypes?**

To address this need, we developed *SLIDE-EX* (SLide-based Inference of DEconvolved gene EXpression), a deep-learning tool trained on deconvolved bulk transcriptomics data that robustly predicts cell type-specific gene expression and cell type abundance from histopathology slides. The prevalence of breast cancer and availability of comprehensive datasets guided our development and optimization of *SLIDE-EX* for this disease. We proceed to provide an overview of *SLIDE-EX*’s framework, the study design, and the cohorts analyzed. We then study the ability to predict cell type-specific data, demonstrating robust predictive performance of the trained model with high gene coverage on an independent, unseen cohort. Next, we applied *SLIDE-EX* to three cohorts of HER2-negative and triple negative breast cancer (TNBC) patients treated with neoadjuvant chemotherapy. Using the predicted cell type-specific gene expression values from these cohorts, we apply our previously published approach, DECODEM^21^, originally developed to predict chemotherapy response from cell-type-specific gene expression profiles deconvolved from measured RNA-seq data, to predict patient response from the *SLIDE-EX* predictions. We show that this framework can accurately predict chemotherapy treatment response using only histopathology slides as the initial input.

## Results

### Study overview

We developed *SLIDE-EX*, a computational pipeline to predict *cell type states* from H&E whole slide images (WSIs) of breast cancer tumors to enable analysis of TME patterns associated with chemotherapy response without requiring resource-intensive molecular profiling. These *cell type states*, comprising cell type-specific gene expression and abundance, were predicted for nine cell types: B-cells, Cancer Associated Fibroblasts (CAFs), Cancer Epithelial (malignant), Endothelial, Myeloid, Normal Epithelial, Plasmablasts, Perivascular-like Cells (PVL), and T-cells.

Using cell type states derived from CODEFACS-analyzed bulk RNA-seq data as ground truth, we developed ten deep learning models: one for cell type abundance and nine for cell type-specific expression. Each model employs image preprocessing with tile extraction, color normalization, and feature extraction using the digital pathology foundational CTransPath^22^ feature extraction, followed by a multi-layer perceptron (MLP) neural network for predicting cell type abundance or TPM normalized gene expression vectors (**Fig. 1a**).

**Fig. 1:**
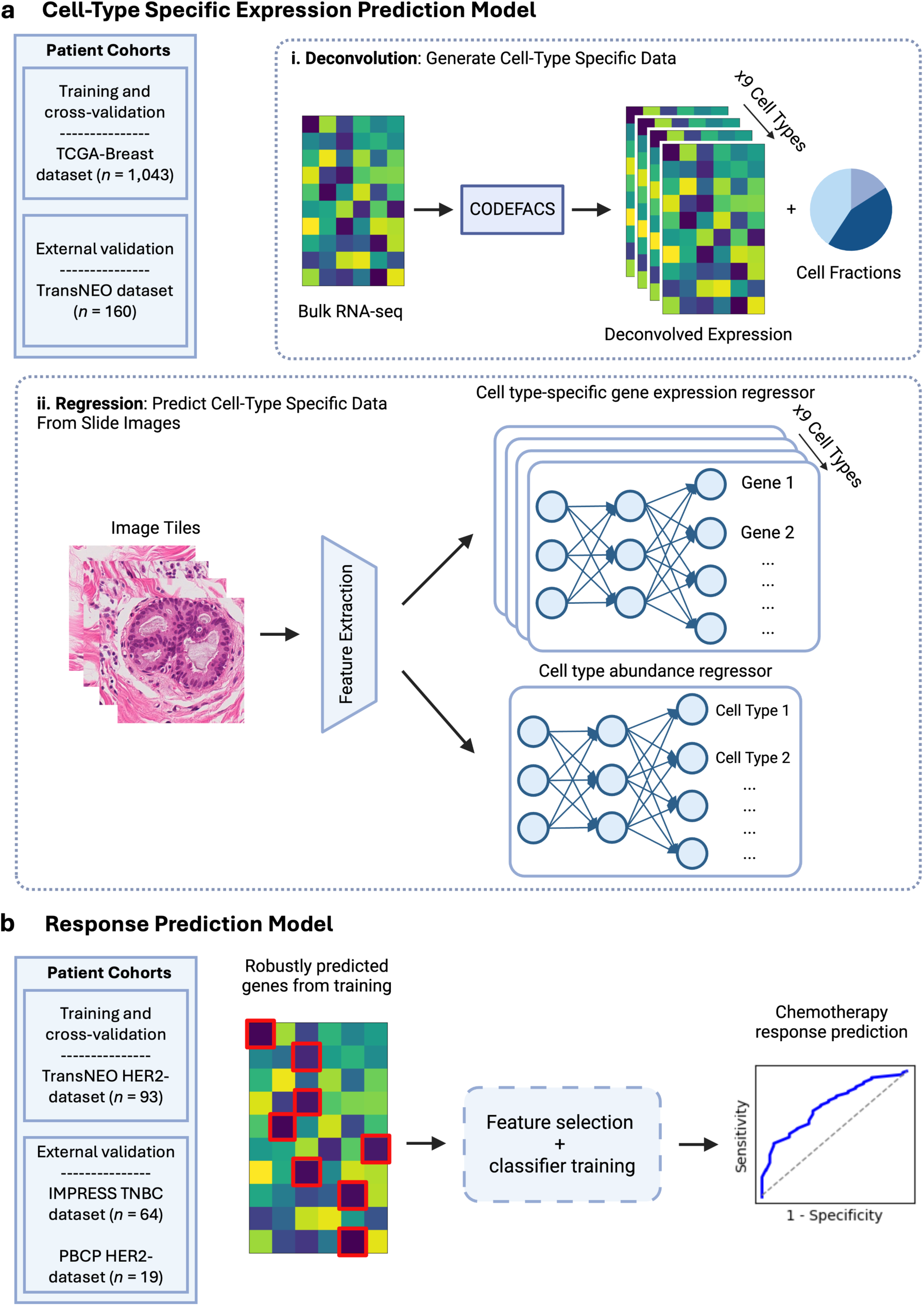
Overview of the datasets and computational workflow of *SLIDE-EX*. **a**, The cell type deconvolution prediction model was trained and evaluated on datasets with paired H&E images and bulk transcriptomics from the TCGA-breast cohort. The TransNEO cohort was used for external validation. (**i**) The bulk transcriptomics data were deconvolved into cell fractions and cell type-specific gene expression using CODEFACS to serve as “ground truth” labels for the machine learning models. (**ii**) Whole slide images (WSIs) were divided into tiles and then color normalized, followed by feature extraction using the CTransPath digital pathology foundation model. The extracted features were then fed into a multi-layer perceptron (MLP) to infer cell type abundance and cell type-specific gene expression for each tile. **b**, DECODEM was used to predict patient response to chemotherapy treatment based on the deconvolved gene expression predictions derived from (**a**). Training and cross-validation were performed on the TransNEO cohort, and the IMPRESS and PBCP cohorts were used for external validation. Only genes that were found to be well-predicted (PCC > 0.4) in the TCGA-breast cohort during training of the cell type-specific gene expression prediction task served as features for the DECODEM response prediction model. The response prediction classifier consists of an ensemble of logistic regression, random forest, and support vector machine models.

We validated these models through cross-validation on 1,043 TCGA breast cancer patients’ WSIs with matched RNA-seq data and assessed generalizability using the TransNEO cohort (*n* = 160).

To evaluate clinical utility, we investigated whether cell type-specific gene expression inferred by *SLIDE-EX* could predict neoadjuvant chemotherapy response using DECODEM, our computational framework for predicting chemotherapy response from deconvolved gene expression profiles (**Fig. 1b**). We trained and cross-validated the response prediction on the HER2-negative subset of the TransNEO cohort (*n* = 93) and validated it on the IMPRESS cohort^23^ (*n* = 64) and the HER2-negative subset of the Personalized Breast Cancer Program (PBCP) cohort (*n* = 19)^3^.

### Histopathology reveals cell type-specific gene expression patterns

As outlined in the overview, *SLIDE-EX* predicts cell type-specific gene expression profiles and cell type abundances directly from histopathology images for each of nine cell type categories. Here we examine the performance of the expression models trained and evaluated using five-fold cross-validation on the TCGA breast cancer cohort. For each cell type, we excluded genes with low expression levels and low variance for each cell type, resulting in predictions for between 11,541 to 16,435 across the nine cell types (**Methods**). Model performance was evaluated using the Pearson correlation coefficient (PCC) between the predicted and cell-type specific expression values of each gene in the test set. Among these, the majority of genes (>97% for all cell types) exhibited a positive correlation in the cross-validation of the TCGA-breast cohort (**Supplementary Table 1, Supplementary Fig. 1a**). Notably, for each of the nine cell types, *SLIDE-EX* predicted at least 1,000 genes with a PCC greater than 0.4, with four cell types—CAFs, Normal Epithelial, B-cells, and Cancer Epithelial—exceeding 2,000 genes above this threshold (**Fig. 2a**). The top-performing cell type predictor was for CAFs, which predicted 2,564 genes with a PCC above 0.4, compared to 3,201 genes predicted by a bulk gene expression predictor trained on the same computational framework. Among the top 1,000 genes with the highest correlations, the averaged median PCC across the nine cell types reached 0.50, ranging from 0.47 (for PVL) to 0.52 (for CAFs, Normal Epithelial, and B-cells) (**Fig. 2b**).

**Fig. 2:**
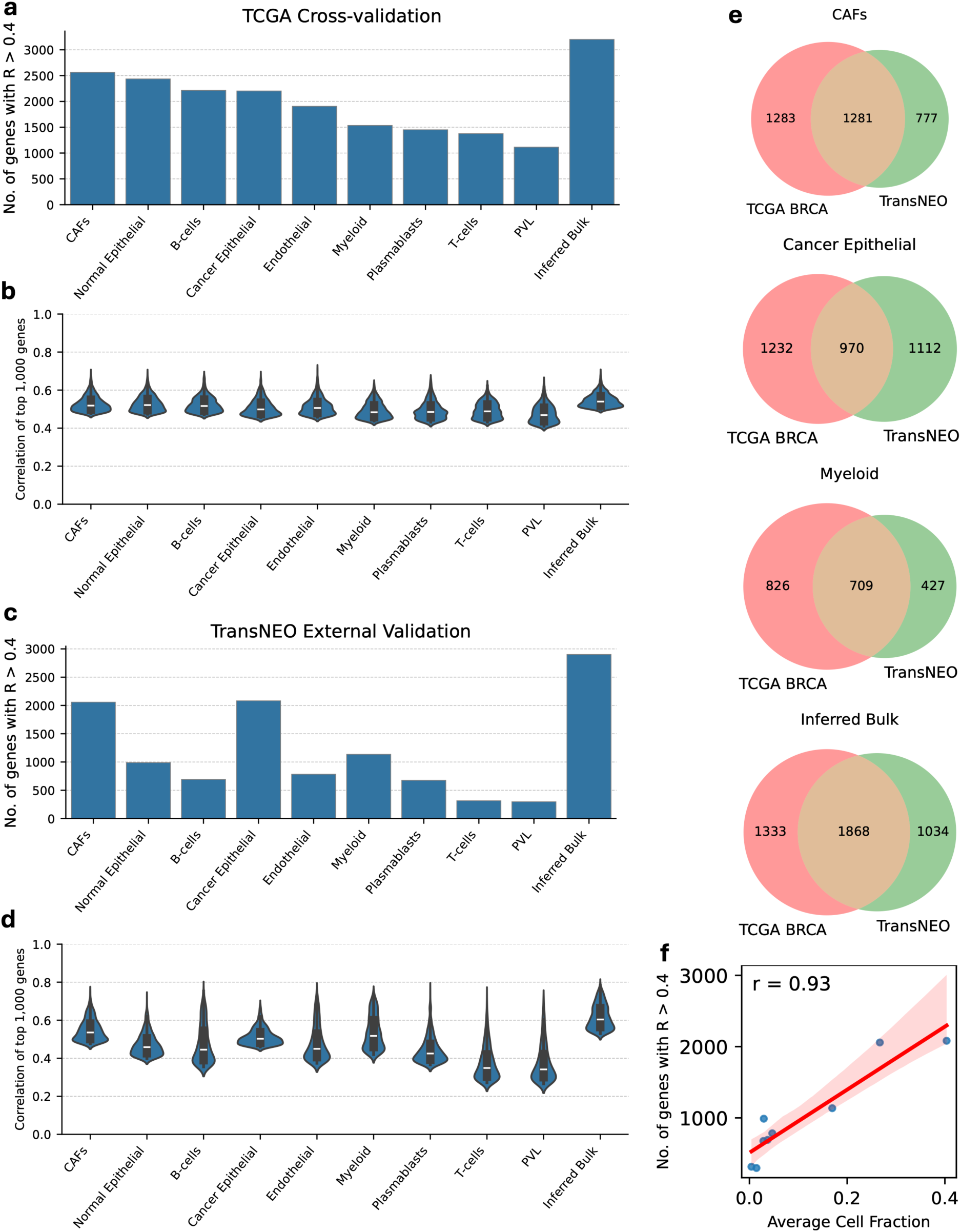
Performance of cell type-specific gene expression prediction from H&E slides. **a-b**, Bar plots of the number of genes with a PCC of >0.4 (**a**) and violin plots depicting the distribution of correlations for the top 1,000 genes with the highest correlations (**b**) in the TCGA-breast cohort (cross-validation). PCCs are calculated between predicted and deconvolved expression values across the cohort for all differentially expressed genes for each subtype. Horizontal lines in violin plots represent median values. **c-d**, Bar plots of the number of genes with a PCC of >0.4 (**c**) and violin plots depicting the distribution of correlations for the top 1,000 genes with the highest correlations (**d**) in the TransNEO cohort (external validation). **e**, Venn diagrams illustrating the overlap between the well-predicted genes (PCC > 0.4) in TCGA-breast (left) and TransNEO (right) for the top 3 performing cell types in the external validation, as well as for the inferred bulk expression. All displayed venn diagrams have hypergeometric *P* values equal to zero. **f**, Scatterplot and linear regression of the number of well-predicted genes (PCC > 0.4) versus the average cell fraction per cell type across the TransNEO external validation cohort (*P* < 1e-3, two-sided t-test).

To assess model generalizability, we applied the cell type-specific gene expression prediction models trained on the TCGA-breast dataset on the TransNEO cohort, an external validation dataset of 160 patients with matched tumor H&E WSIs and gene expression data. Notably, *SLIDE-EX* demonstrated robust performance despite substantial differences in sample preparation—with TCGA samples derived from formalin-fixed, paraffin-embedded (FFPE) surgical resections and TransNEO samples from fresh-frozen needle biopsies (**Supplementary Fig. 2**). Across all cell types, the averaged median PCC for the top 1,000 genes was 0.45, ranging from 0.34 for PVL to 0.54 for CAFs (**Fig. 2d**). Furthermore, the CAF and cancer epithelial cell predictors each achieved over 2,000 well-predicted genes (PCC > 0.4) (**Fig. 2c**).

We observed a significant overlap between well-predicted genes in the training and external validation cohorts for each cell type (*P* < 1e-80, two-sided hypergeometric test), demonstrating the consistency of our model. Particularly, of the 2,058 well-predicted genes for CAFs in the TransNEO cohort, 62% (1,281 genes) overlapped with those in the TCGA-breast cohort (**Fig. 2e**). Similarly, among the 2,082 well-predicted genes for Cancer Epithelial cells in the TransNEO cohort, 47% (970 genes) overlapped with those in the TCGA-breast cohort, and of the 1,136 well-predicted genes for Myeloid cells in the TransNEO cohort, 62% (709 genes) overlapped with those in the TCGA-breast cohort (**Fig. 2e, middle**). The cell type-specific models showed similar consistency to the bulk gene expression model, where 64% (1,868) of the 2,902 well-predicted TransNEO genes overlapped with the TCGA-breast cohort (**Fig. 2e**).

In addition, we observed a strong correlation between cell type abundance and *SLIDE-EX*’s prediction accuracy in the TransNEO cohort (**Fig. 2f**), with performance decreasing for less abundant cell types—potentially reflecting an increased signal-to-noise ratio in more abundant cell types—a trend also reported in the performance of CODEFACS itself^13^.

### Histopathology images reveal cellular composition of the TME

In parallel with predicting cell type-specific gene expression, we trained a model to predict the cell type fractions, or abundances, for the same nine cell types directly from histopathology image features in the TCGA-breast cohort. Using RNA-seq-derived cell type abundances as ground truth, we assessed model performance through PCCs between predicted and deconvolved cell fractions across all nine cell types in both training and validation datasets. During training and cross-validation, all cell types yielded a positive correlation (*P* < 0.05 for all cell types, two-sided t-test) (**Supplementary Table 2**). Additionally, three out of the nine cell types had a PCC exceeding 0.4, specifically CAFs (PCC = 0.56), Myeloid (PCC = 0.53), and Cancer Epithelial (PCC = 0.46) (**Fig. 3a**).To further validate the cell fraction predictor’s performance and demonstrate its generalizability, we again tested the model on the corresponding fresh-frozen H&E slides and deconvolved cell fractions of the TransNEO cohort. The top three performing cell types were the same in the independent validation, displaying PCCs above or around 0.4, and all cell types maintained a positive correlation. This suggests that our model can remain robust and generalizable across patient cohorts and tissue preparation methods. Surprisingly, the prediction correlation for Myeloid cells noticeably increased in the external validation compared to cross-validation, with a PCC of 0.71. The other two top cell types maintained similar performances to cross-validation (CAFs, PCC = 0.58; Cancer Epithelial, PCC = 0.39; **Fig. 3b**).

**Fig. 3:**
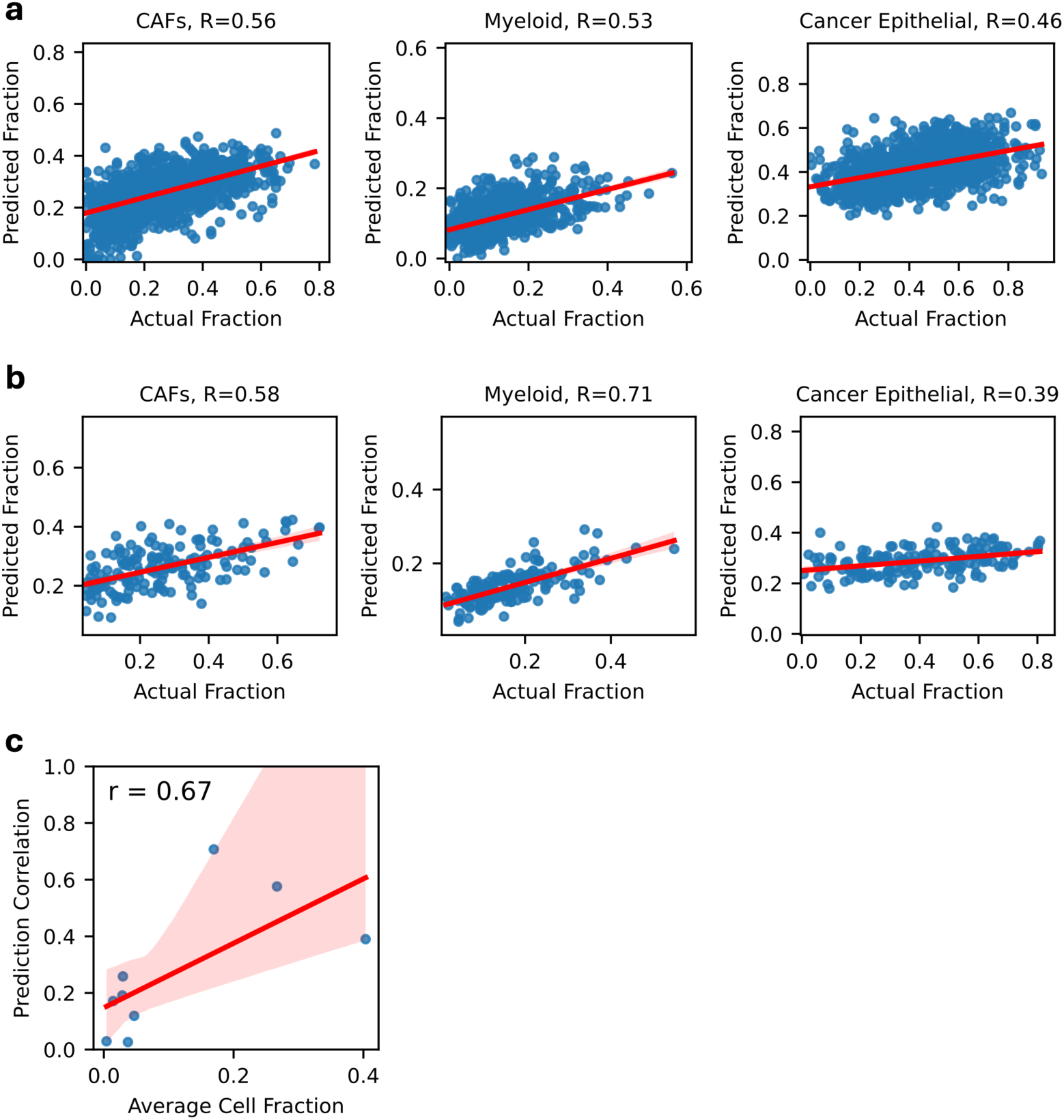
Performance of cell-type abundance prediction from H&E slides. **a-b**, Scatterplots of the predicted versus deconvolved fractions for the top 3 cell types with the highest PCCs in the TCGA-breast cross-validation cohort **(a)** and for the same 3 cell types in the TransNEO external validation cohort **(b). c**, Scatterplot and linear regression of the PCC between the predicted and deconvolved cell fractions versus the average cell fraction per cell type across the TransNEO external validation cohort (*P* < 0.05, two-sided t-test).

As with cell type-specific expression, *SLIDE-EX*’s ability to infer abundances is correlated with cell type abundances (**Fig. 3c**), highlighting that *SLIDE-EX* inferences for cell type fractions from WSIs are more robust for abundant cell types.

### Clinical subtype influences cell type state prediction performance

We sought to investigate whether gene expression prediction performance is affected by the clinical subtype of the breast cancer tumor. We stratified the patients in the TCGA-breast cohort, this time by clinical subtype, and analyzed the distribution of prediction correlations among patients sharing the same clinical subtype. The three subtypes studied were TNBC (*n* = 120), HR positive (*n* = 488), and HER2 positive (*n* = 107). We observed that the median correlation between the predicted and deconvolved values was lower for TNBC patients than for HR positive and HER2 positive patients across all cell types (**Fig. 4a**), despite TNBC patients representing an intermediate-sized cohort between the larger HR positive and smaller HER2 positive groups. The difference in distributions was statistically significant for all nine cell types (paired Wilcoxon signed-rank test *P* < 0.001). This trend of lower TNBC predictive performance persisted when looking at the number of well-predicted genes by subtype, with the TNBC subset having fewer well-predicted genes than both the HR positive and HER2 positive subsets (**Fig. 4b**), and when looking at the distributions of the top 1,000 well-predicted genes in each subtype (**Supplementary Fig. 3**), suggesting that the unique morphological features of TNBC make it a challenging population for histopathology-based biomarker prediction tools.

**Fig. 4:**
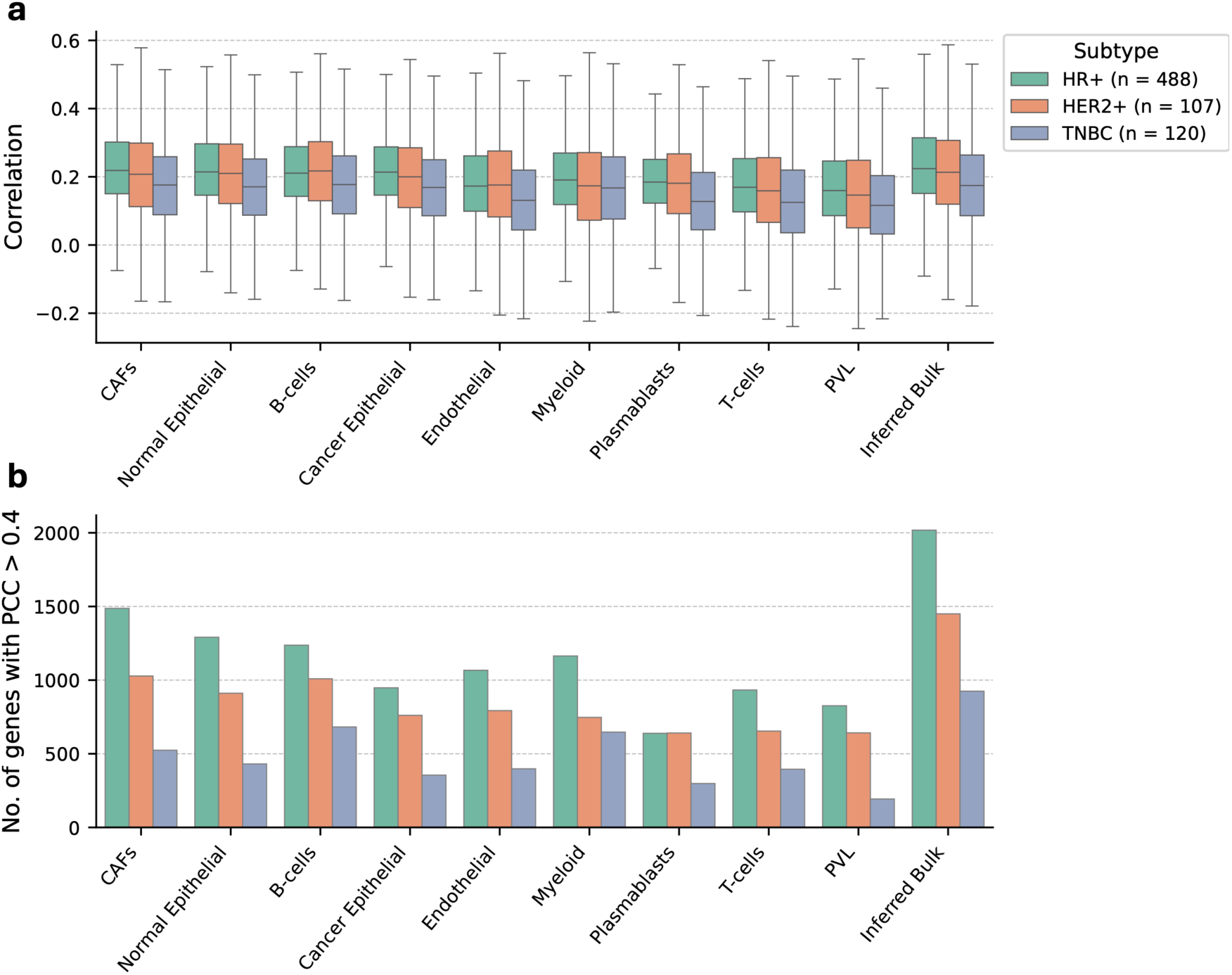
Performance of cell type-specific gene expression prediction across breast cancer subtypes. **a**, Box plots of the PCCs for the predicted genes across the breast cancer clinical subtypes in the TCGA-breast cohort. **b**, Bar plots of the number of genes with a PCC of >0.4 across the breast cancer subtypes in the TCGA-breast cohort.

### Robustly predicted genes reflect key biological functions of their respective cell types

We next explored whether genes that are robustly predicted (specifically having a PCC of >0.4 between predicted and deconvolved expression in both the TCGA-breast and TransNEO cohorts) have known biological relevance to cancer or to tumor immune microenvironment processes. To this end, we performed a Gene Ontology (GO) enrichment analysis on gene sets of the Biological Process aspect. **Fig. 5** summarizes the enrichment analysis results for the nine cell types of interest. Reassuringly, this analysis highlights the specific enrichment of cell cycle and DNA replication pathways among cancer cells. These are classical cancer hallmark pathways that are often disrupted in breast cancer^24–26^ and promote tumor proliferation, making them important prognostic markers and therapeutic targets^27–30^. Immune cells, on the other hand, exhibited enrichment in processes involved in immune response regulation, such as the activation and proliferation of T-cells, B-cells, and lymphocytes, and positive regulation of cytokine production. CAFs showed enrichment in pathways related to extracellular matrix organization and regulation of cell migration. These cell type-specific enrichment patterns demonstrate that *SLIDE-EX* successfully captures genes central to each cell type’s biological function.

**Fig. 5:**
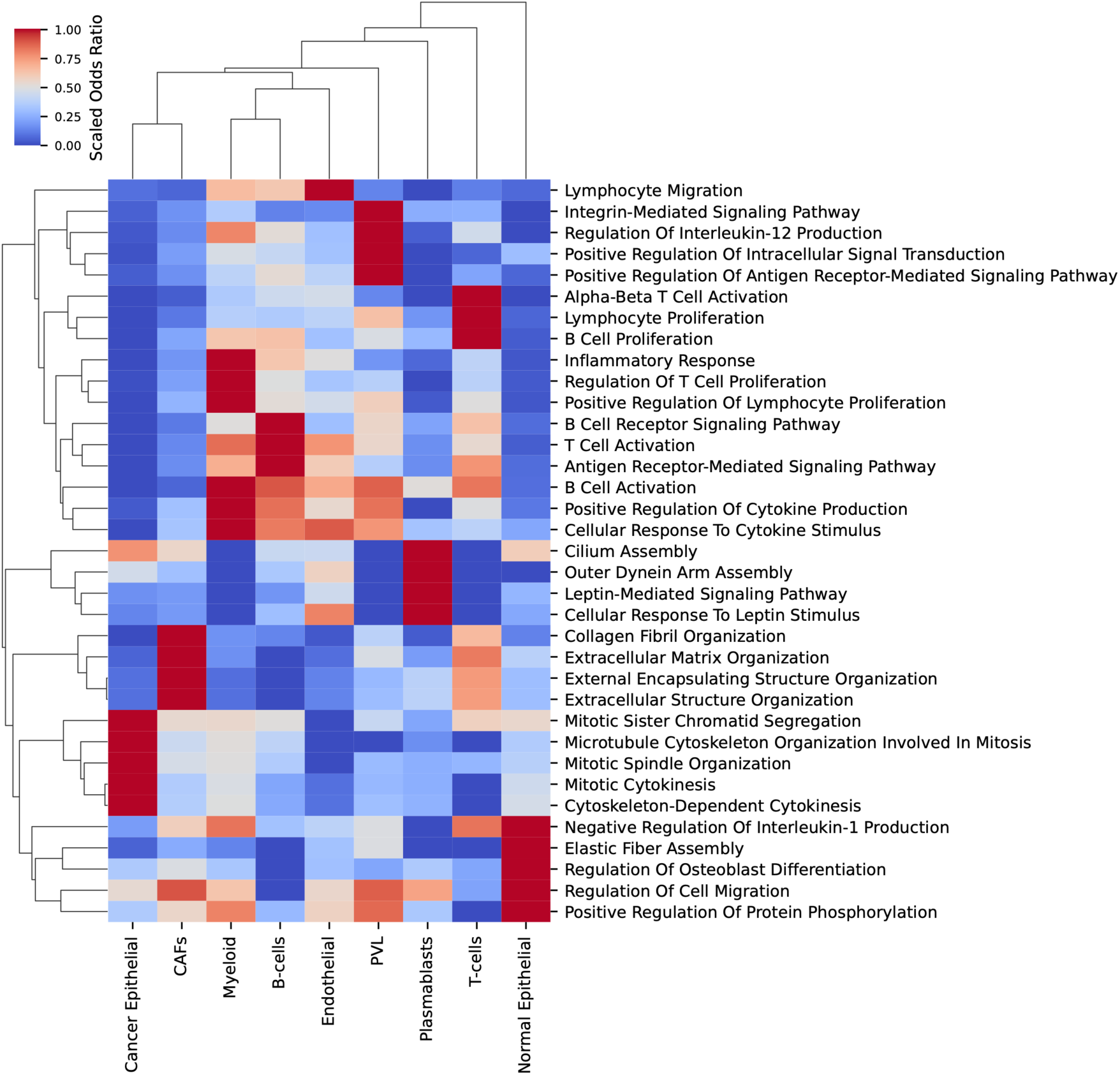
GO pathway enrichment analysis on the intersection of well-predicted genes (PCC > 0.4) in both cross-validation and external validation. The pathways displayed are the union of the top 5 most significantly enriched pathways for each cell type (hypergeometric test). Values (0 to 1) denote the odds ratios scaled per cell type (column).

### The predicted cell type-specific gene expression predicts chemotherapy response

After establishing the predictive ability of these cell type-specific gene expression models, we sought to demonstrate their clinical utility by evaluating their ability to predict patient response to neoadjuvant chemotherapy. For this analysis, we utilized a subset of the TransNEO cohort, comprising 93 HER2-negative patients who received neoadjuvant chemotherapy and had available WSIs with corresponding response data. From these WSIs, we inferred cell type-specific gene expression for each of the nine cell types. Using the inferred gene expression from each cell type as features, we utilized DECODEM to develop cell type-specific machine learning classifiers to predict complete pathological response (pCR; *n* = 21). Each classifier was evaluated through cross-validation.

To benchmark our approach, we developed two baseline predictors: one utilizing inferred bulk expression values for classification, termed *inferred bulk*, and a second that classified patient response directly from image features by using the same computational pipeline as our expression predictors but replacing the regression head with a classification head. We termed this second approach *the direct supervised model*.

In cross-validation on the TransNEO cohort, all nine cell type-specific models outperformed the direct model in terms of both area under the receiver operating characteristic curve (AUC) and average precision (AP; equivalent to the area under the precision-recall curve), and all exhibited similar performance to the bulk predictor (**Fig. 6a-b**; left). In addition, we observed a positive correlation between the prediction AUC of a cell-type specific model and the number of well-predicted genes (PCC > 0.4) for that cell type in the initial TCGA-breast cohort (**Supplementary Fig. 4a-b**).

**Fig. 6:**
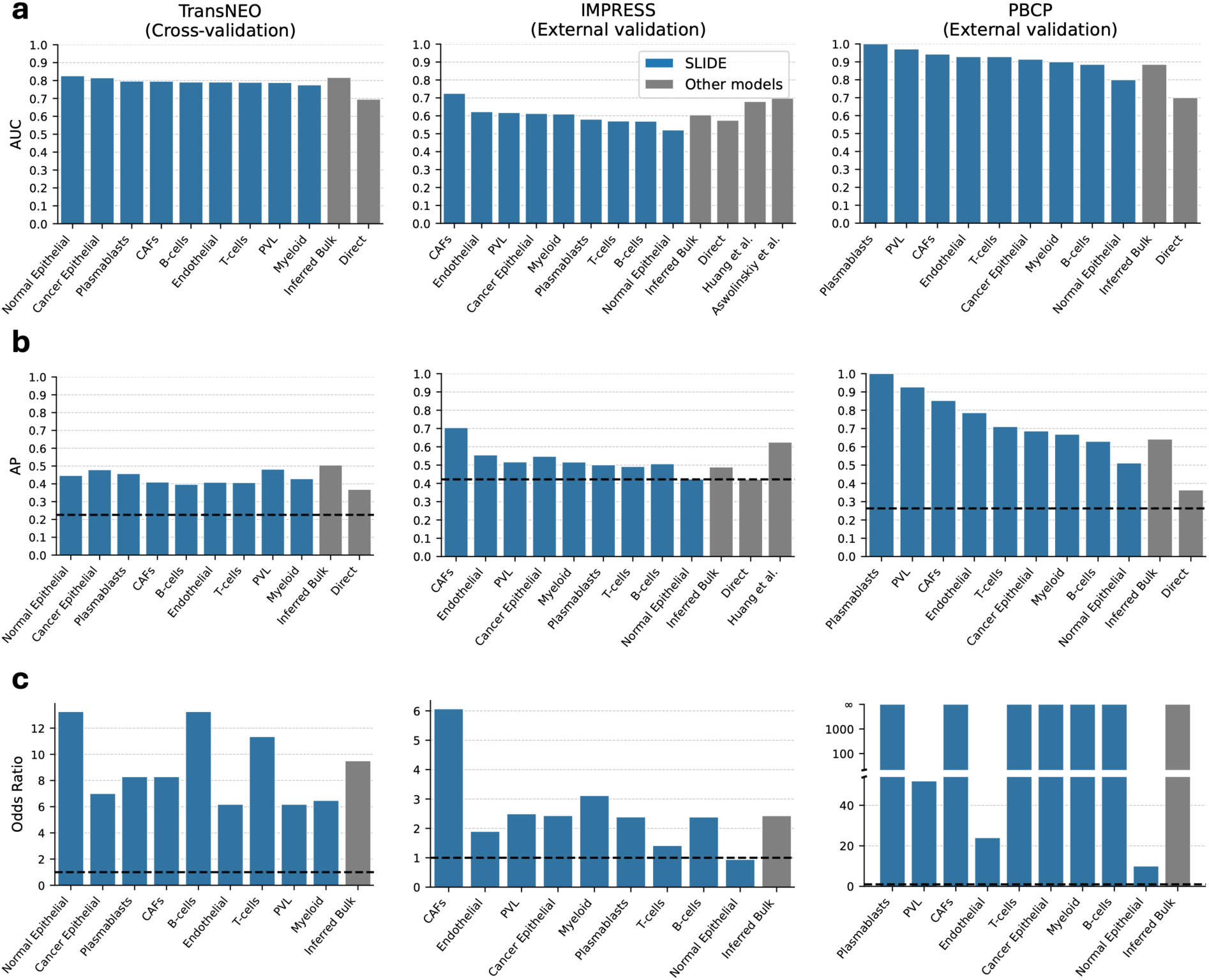
Predicting chemotherapy treatment response from H&E slides. **a**, Bar plots of AUC values of response prediction for nine cell type-specific models compared to inferred bulk and direct predictors across three cohorts, with cell types ranked by descending values in cross-validation. Results shown for TransNEO HER2-cross-validation (*n* = 93; left), IMPRESS external validation (*n* = 64; middle), and PBCP external validation (*n* = 19; right). **b**, Average precision (AP) values for all models across three cohorts, with dashed lines indicating each cohort’s overall response rate (ORR); An AP higher than the ORR demonstrates better-than-chance accuracy. Results shown for TransNEO HER2-cross-validation (*n* = 93; left), IMPRESS external validation (*n* = 64; middle), and PBCP external validation (*n* = 19; right). **c**, Odds ratio values for all models across three cohorts, with dashed lines at OR = 1 indicating the threshold for positive association. Results shown for TransNEO HER2-cross-validation (*n* = 93; left), IMPRESS external validation (*n* = 64; middle), and PBCP external validation (*n* = 19; right).

To assess the generalizability and robustness of our models, we validated the predictive power of all cell type-specific models in external validation on two independent cohorts. We first used the IMPRESS cohort, comprising 64 TNBC patients who received neoadjuvant chemotherapy (*n* = 27 pCR) (**Fig. 6a-b**; middle). This validation was particularly challenging as models trained on TransNEO’s mixed subtypes were evaluated on IMPRESS’s exclusively TNBC cases. Notably, four cell type models achieve a higher test AUC than the bulk expression model. In particular, the CAF classifier reached an AUC of 0.72, compared to 0.60 for the bulk model (**Fig. 6a**; middle). For prediction performance assessed by AP, all cell type models except normal epithelial exceed the overall response rate (ORR), reaffirming the predictive power of this pipeline (**Fig. 6b**; middle; dotted line). Importantly, our CAF-based model outperformed existing H&E-based methods for predicting chemotherapy response in breast cancer on the same challenging IMPRESS dataset, which reported AUCs of 0.68 (Huang et al.^23^; cross-validation) and 0.70 (Aswolinskiy et al.^31^; external validation). This demonstrates that cell type-specific transcriptional information derived from histopathology images can provide superior predictive power compared to conventional image-based approaches.

We then validated our models on a subset of the PBCP cohort, comprising 19 HER2-negative patients who received neoadjuvant chemotherapy (*n* = 5 pCR) (**Fig. 6**; right). This population closely resembled the HER2-negative patients we examined in the TransNEO cohort cross-validation. Notably, the Plasmablast classifier achieved perfect discrimination on the PBCP cohort, with both AUC and AP reaching 1.0. This finding aligns with clinical evidence that high densities of tumor-infiltrating plasmablasts and plasma cells, quantified through associated marker expression, predict chemotherapy response in breast cancer^32–34^. Furthermore, 8 of the 9 cell type-specific models demonstrated test AUCs exceeding or equal to the bulk model’s AUC of 0.89, and all cell type models surpassed the direct model (**Fig. 6a**; right). The APs attained by all cell type models substantially exceeded the ORR for the PBCP cohort, indicating robust predictive performance (**Fig. 6b**; right; dotted line).

To assess potential clinical applicability, we evaluated our models using the odds ratio (OR) of response. The OR quantifies the likelihood of response in patients predicted to respond compared to those predicted not to respond. Across all cell type-specific models, except for normal epithelial cells in IMPRESS, the odds were higher for predicted responders (OR > 1) (**Fig. 6c**). Notably, the CAF model, the top performing cell type in IMPRESS external validation, achieved an OR of 6.1, further indicating that patients classified as responders by our WSI-inferred cell type-specific expression models were more likely to achieve complete pathological response to chemotherapy, even in this challenging all-TNBC cohort (**Fig. 6c**; middle). In the PBCP cohort, 6 cell-type specific models, as well as the bulk model, achieve perfect separation, indicated by an OR of infinity (**Fig. 6c**; right).

## Discussion

This study presents *SLIDE-EX*, a deep-learning framework designed to predict cell type states directly from histopathology images. We first demonstrate its ability to predict deconvolved gene expression profiles and cellular fractions through cross-validation and external validation on tumor samples of various breast tumor subtypes. Furthermore, we show that the cell type-specific gene expression profiles generated by *SLIDE-EX* predict chemotherapy response in three cohorts using only H&E images, outperforming existing approaches.

We also investigated how model performance varied across different breast cancer subtypes, where we found that predicting cell type-specific gene expression in TNBC cases showed lower accuracy compared to other subtypes. Notably, the robustly predicted genes for each cell type were enriched in Gene Ontology (GO) terms characteristic of the biological processes of the corresponding cell type, highlighting the biological relevance of *SLIDE-EX*’s predictions.

While our approach shows strong potential in predicting cell type-specific gene expression profiles directly from histopathology slide images, it has certain limitations. Notably, prediction performance was lower for less abundant cell types, likely due to their reduced representation in the images. Their limited presence may result in less distinguishable histological features, making it more challenging for the model to capture meaningful patterns associated with these cells. Additionally, our approach depends on the accuracy of cell type deconvolution from bulk RNA-seq data. We employed CODEFACS for deconvolution, and any limitations inherent to this method—such as variability in RNA-seq processing, reliance on well-defined molecular signatures, and a similar decrease in performance for less abundant cell types—can impact the downstream performance of *SLIDE-EX*. Ultimately, *SLIDE-EX* provides an approximation of cellular states that can offer valuable biological insights and even predict treatment response in scenarios where resource-intensive single-cell or bulk RNA-seq data are unavailable. However, the insights derived from *SLIDE-EX* should be validated using single and spatial sequencing approaches, underscoring the continued importance of gene expression assays for precise prognostic evaluation.

Overall, *SLIDE-EX* provides a cost-effective approach for predicting cell type-specific gene expression and cell type abundances, facilitating large-scale investigations that were previously constrained by limited access to transcriptomic assays. By enabling the exploration of gene expression patterns in patient cohorts lacking matched transcriptomic data, *SLIDE-EX* opens new avenues for uncovering key insights into the tumor microenvironment and its underlying molecular landscape in potentially many cancer types. This advancement lays the groundwork for more personalized treatment strategies and, ultimately, improved patient outcomes.

## Methods

### Data collection

The datasets used in this study come from publicly available resources. Details are described below.

TCGA histological images and their corresponding gene expression profiles were downloaded from the Genomic Data Commons Data Portal (https://portal.gdc.cancer.gov). Only diagnostic slides from primary tumors were selected, making a total of 1,106 FFPE slides from 1,043 patients with breast cancer.

The TransNEO-breast dataset consists of FF slides and their corresponding gene expression profiles from 160 patients with breast cancer. Full details of the RNA library preparation and sequencing protocols, as well as digitization of slides, have been previously described^3^.

The IMPRESS-TNBC dataset consists of histopathology slides from 64 TNBC patients treated with neoadjuvant chemotherapy and subsequent surgical excision^23^.

### *SLIDE-EX* computational framework

Our model architecture consists of four main components (**Fig. 1a**).

### Deconvolution with CODEFACS

The first step involves employing CODEFACS, a cellular deconvolution tool recently developed in our lab^13^. CODEFACS characterizes the TME by reconstructing the cell type-specific expression in each sample from the input measured bulk expression, in transcripts per million (TPM), utilizing the corresponding cell type abundance or a cell-type-specific signature profile as another input. The output includes the cell type-specific expression profiles in TPM and the corresponding cell type abundances in each cell type across all samples.

In a previous work, we derived a cell type signature matrix of 1,400 genes representing nine cell types from a single-cell RNA-seq (scRNA-seq) cohort from Wu et al.^35^. We then deconvolved the bulk expression profiles with this signature using CODEFACS into cell-type-specific expression profiles, encompassing nine cell types: B-cells, Cancer-associated Fibroblasts (CAFs), Cancer Epithelial (malignant), Endothelial, Myeloid, Normal Epithelial, Plasmablasts, Perivascular-like Cells (PVL), and T-cells.

### Image processing

The preprocessing phase begins by dividing whole-slide images into non-overlapping small images known as tiles. Each tile size was standardized to 512 x 512 pixels across all cases. A magnification level of 20× was chosen for analysis. Tiles that predominantly comprised more than half of the background were omitted. Consequently, each WSI could be represented by several thousand tiles, depending on the slide’s dimensions. Color normalization techniques were employed to mitigate staining discrepancies across slides.

### Feature extraction

Image tiles were processed using CTransPath^22^, a foundational digital pathology model trained through self-supervision on histopathology images. Each tile is ultimately represented by a 768-dimensional CTransPath feature vector.

### Multilayer perceptron regression

This component builds a predictive model linking CTransPath features to whole-genome gene expression at a cell type-specific level. The model architecture consists of three layers: (1) an input layer with 768 nodes, corresponding to the size of the CTransPath feature vector; (2) a hidden layer with 768 nodes; and (3) an output layer with one node per gene.

In order to facilitate convergence of our deep learning model, we excluded genes with low expression levels and low variance for each cell type, as well as genes that were not common to both the TCGA and TransNEO bulk expression profiles. Specifically, genes with non-zero expression in less than 50% of the cohort were excluded. In addition, genes with an expression level variance across the cohort of less than 1e-5 were excluded. This threshold was determined empirically as the smallest variance that did not lead to interference with model convergence. Expression values were converted from TPM to logTPM, calculated as log(TPM + 1), before training. Since the deconvolved expression profiles varied from cell type to cell type, the number of genes predicted ranged from 11,541 to 16,435.

Because the training data consist of gene expression at the slide level (that is, bulk gene expression, as opposed to at spatial resolution), we averaged our per-tile predictions to obtain a mean value at the slide level. Alternatively, when predicting cell type abundance, the third and final layer instead has 9 nodes, one for each cell type. To ensure that the 9 cell type abundance predictions are all non-negative and sum up to 1, the outputs are normalized by first subtracting the minimum value from the output vector if the minimum is negative, and then dividing each value by the sum of the output values.

### Model training and evaluation

To evaluate the performance of our models, we applied 5 × 5 nested cross-validation. For each outer loop, we split the entire patient population (of each cohort) into training (80%) and held-out test (20%) sets. We further split the training set into internal training and evaluation sets according to fivefold cross-validation. The models were trained and evaluated independently with each pair of training/validation sets. Averaging the predictions from the five different models represents our final prediction for each single gene and cell type on each held-out test set. We repeated this procedure five times across the five held-out test sets, making a total of 25 trained models. These models trained with the TCGA cohort were used to predict the expression of each gene and the abundance of each cell type in the external TransNEO cohort by computing the mean over the predicted values of all models.

The models were trained using stochastic gradient descent with a minibatch size of 32. For optimization, we employed the Adam optimizer with a learning rate of 0.0001. The mean squared error between predicted and deconvolved values was used as the loss function. To reduce the risk of overfitting, a dropout rate of 20% was applied to the initial layer. Each training round was conducted over 500 epochs. We also implemented early stopping to enhance computational efficiency and further prevent overfitting. The training was stopped if no improvement was observed in the mean correlation between predicted and deconvolved values on the validation set after 50 continuous epochs.

### Analysis of predicted cell type-specific gene expression

For downstream analysis of the predicted cell type-specific gene expression, we utilized averaged values from the 25 trained models in cross-validation as mentioned above.

Data on the cancer stage and subtype of each patient was obtained from the Genomic Data Commons Data Portal.

Gene Ontology (GO) enrichment analysis was performed using the Python package GSEAPY^36^ with the enrichr function for overrepresentation analysis. Gene sets were from the GO Biological Process aspect (The specific version is the ‘GO_Biological_Process_2023’ gene-set library from Enrichr)^37–39^. The cluster heatmap was plotted for the union of the top 5 enriched pathways of each cell type (**Fig. 5**).

### Chemotherapy response prediction

DECODEM was used to predict patient response to chemotherapy treatment from the *SLIDE-EX* expression predictions. DECODEM takes in deconvolved gene expression profiles as input and trains an ensemble machine learning classifier as described in our previous work^21^. The ensemble consists of regularized logistic regression, random forest, and support vector machine models.

In the response prediction workflow, cell type-specific gene expression values were inferred via *SLIDE-EX* for each patient based on their corresponding H&E slide images. Then, these output expression predictions were fed into DECODEM. Of note, we included only genes that demonstrated robust prediction performance (PCC > 0.4) in the TCGA-breast cohort in cross-validation. This feature selection step was based on our hypothesis that well-predicted genes would provide stronger discriminating signals.

### Direct supervised model

The direct (end-to-end) supervised model was designed to classify responders and non-responders directly from their slides without the intermediate step of cell type-specific gene expression prediction. To this end, we applied the same computational deep-learning framework that was used for the prediction of gene expression, except that the MLP regression component was replaced by an MLP classification component. The same nested cross-validation approach was used to train and evaluate this model, except the TransNEO cohort was the cross-validation dataset and the IMPRESS cohort was the external validation dataset. For each model, slide-level prediction was computed by averaging tile-level predictions within that slide. A threshold of 0.5 was used to determine responder vs. non-responder from the model output.

### Implementation details

Analysis in this study was mainly performed in Python 3.9.7 with libraries including Numpy 1.20.3, Pandas 1.3.4, Scikit-learn 1.2.2 and Matplotlib 3.4.3. Image processing including tile partitioning and color normalization was conducted with OpenSlide 1.1.2, OpenCV 4.5.4 and PIL 8.4.0. The feature extraction, MLP regression, and MLP classification components were implemented using PyTorch 1.12.0.

