## Supplementary Information for "Deep learning inference of cell type-specific gene expression from breast tumor histopathology"

### Supplementary Tables

| Cell Type | Percent of genes with positive correlation |
|---|---|
| Normal Epithelial | 99.79 |
| B-cells | 99.79 |
| CAFs | 99.64 |
| Cancer Epithelial | 99.53 |
| Plasmablasts | 99.49 |
| Myeloid | 99.00 |
| PVL | 98.31 |
| Endothelial | 98.29 |
| T-cells | 97.98 |

**Supplementary Table 1: Percent of genes with a positive PCC for each of the nine cell types in the TCGA-breast cross-validation cohort.**

| Cell Type | Correlation on Cross-validation | Correlation on External Validation |
|---|---|---|
| CAFs | 0.5594739457 | 0.5760176808 |
| Myeloid | 0.5311360156 | 0.7070633196 |
| Cancer Epithelial | 0.4579560635 | 0.3899380889 |
| Endothelial | 0.3679009611 | 0.1191713522 |
| B-cells | 0.3511271203 | 0.02657731463 |
| Plasmablasts | 0.3226170017 | 0.1911109745 |
| Normal Epithelial | 0.3158136056 | 0.2584835173 |
| T-cells | 0.2127615023 | 0.02925149912 |
| PVL | 0.06571688573 | 0.1715426586 |

**Supplementary Table 2: Pearson correlation coefficient between the predicted and deconvolved cell fractions for each of the nine cell types.** Cross-validation was performed on the TCGA-breast cohort, and external validation was performed on the TransNEO cohort. Cell types sorted by decreasing PCC on cross-validation.

### Supplementary Figures

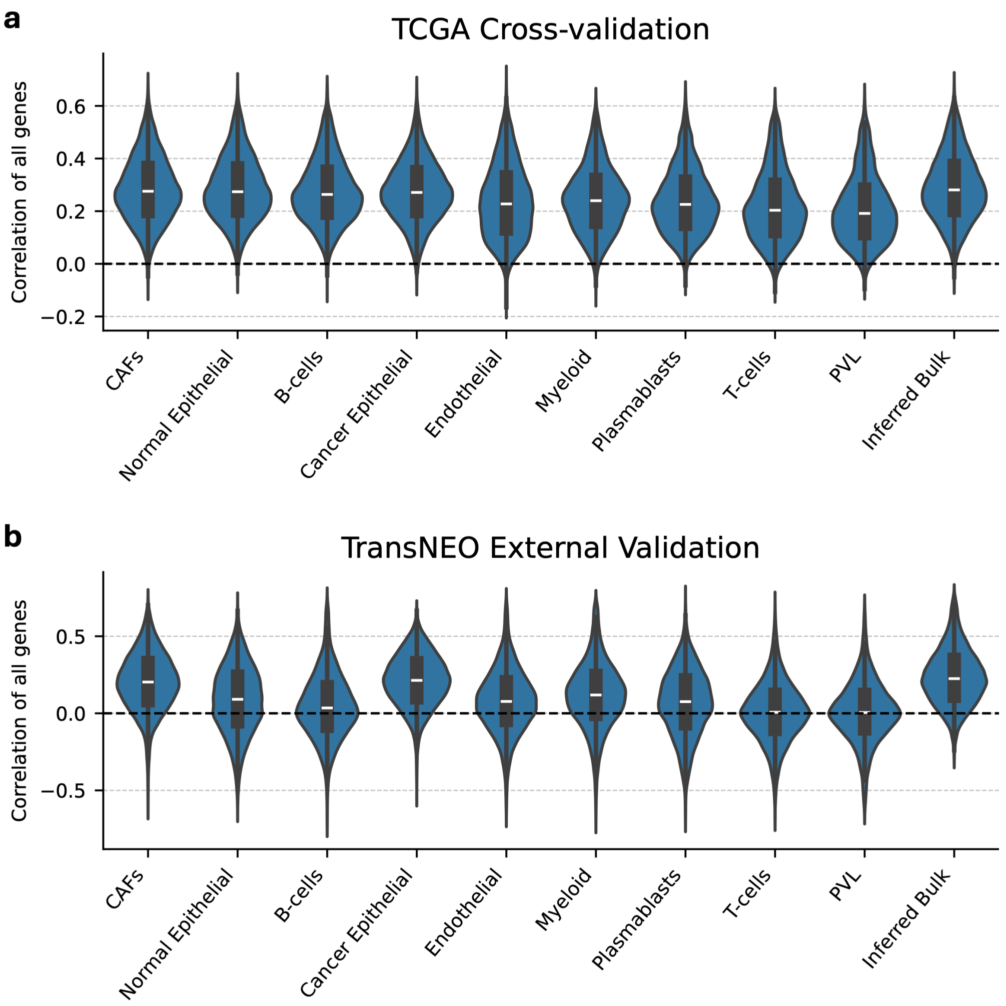


**Supplementary Fig. 1: Performance of cell type-specific gene expression prediction across all genes. a-b,** Violin plots depicting the distribution of correlations for all predicted genes (**a**) in the TCGA-breast cross-validation cohort and (**b**) in the TransNEO external validation cohort.

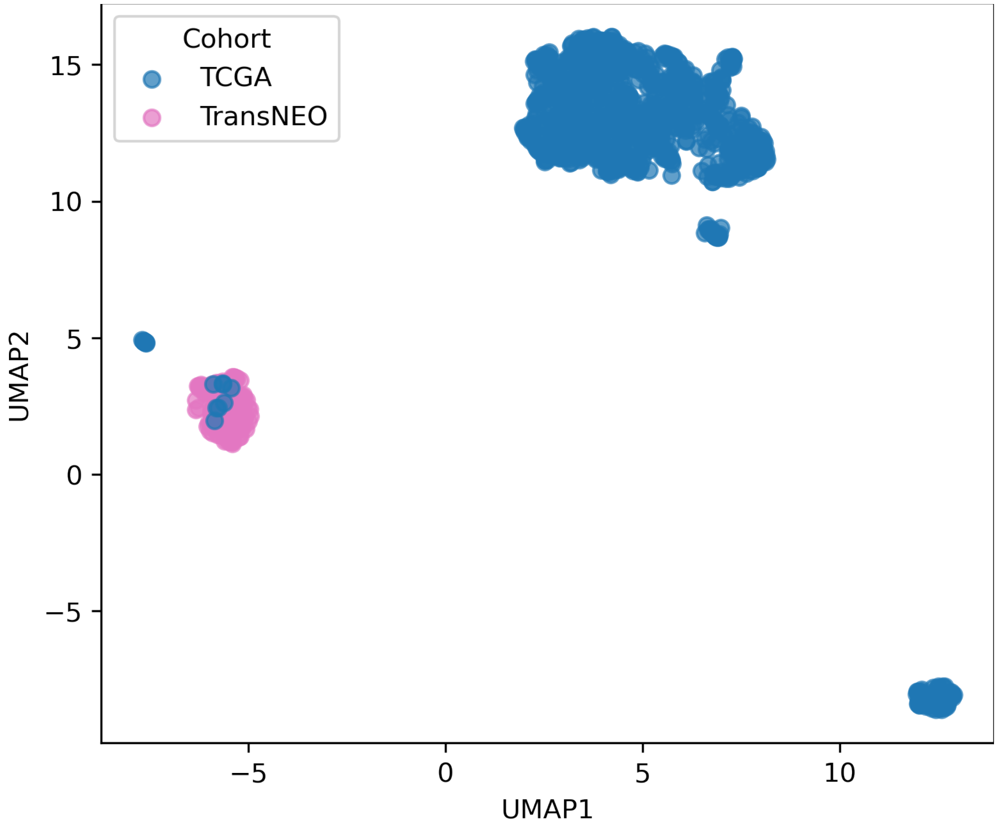


**Supplementary Fig. 2: UMAP projection of WSIs from TCGA and TransNEO cohorts in CTransPath embedding space.** For each WSI, the per-tile feature vectors from CTransPath were averaged to obtain a single mean embedding vector that represents the entire slide. The mean embedding vectors were then projected into UMAP space.

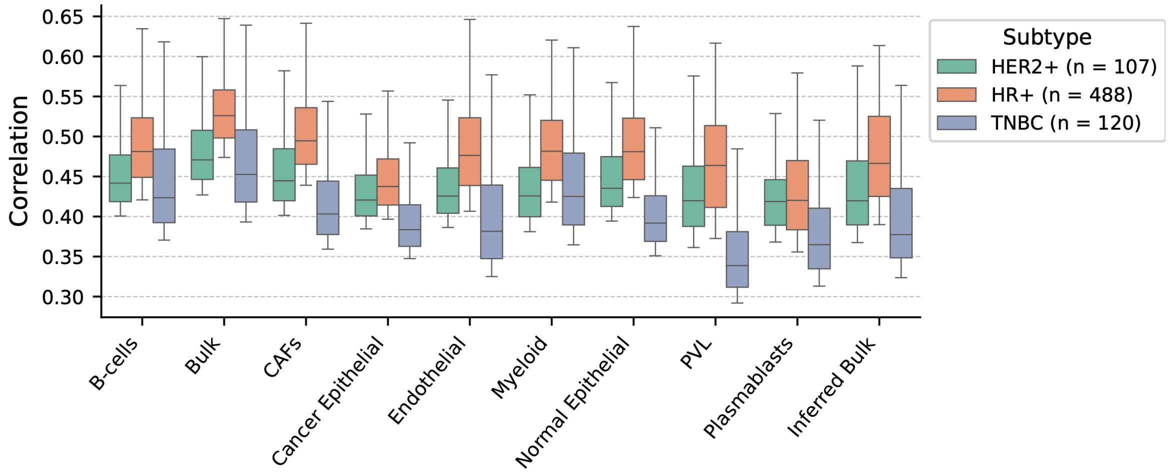

**Supplementary Fig. 3: Top performing genes in cell type-specific gene expression prediction across breast cancer subtypes.** Box plots of the top 1,000 PCCs for the best predicted genes across the breast cancer clinical subtypes in the TCGA-breast cohort.

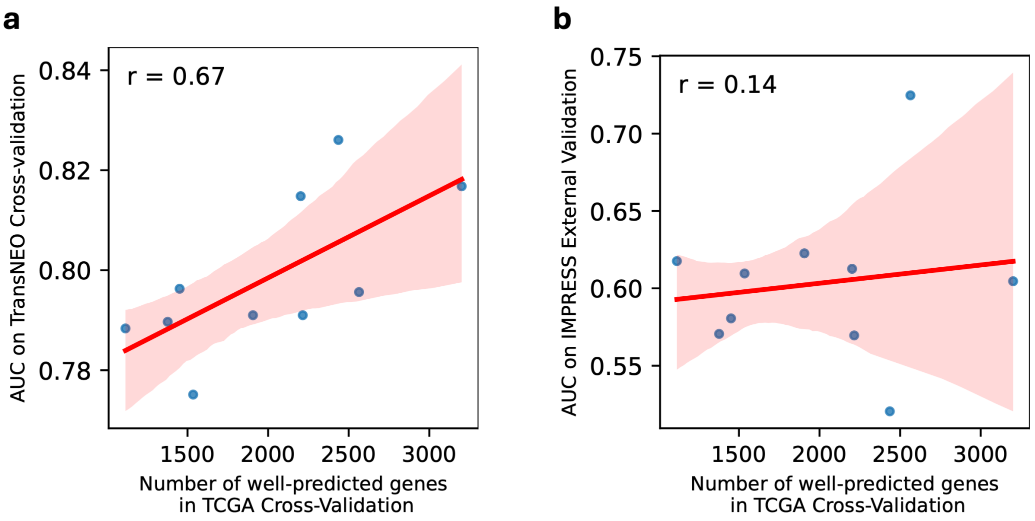


**Supplementary Fig. 4: Correlation between number of well-predicted genes and chemotherapy treatment response prediction performance. a-b**, Scatterplot and linear regression of the AUCs of each cell-type specific treatment response predictor in TransNEO cross-validation cohort (**a**) and IMPRESS external validation cohort (**b**) versus the number of well-predicted genes (PCC > 0.4) for that cell type in the initial TCGA-breast cohort.
